# Global Proteomic Analyses of Macrophage Response to *Mycobacterium tuberculosis* Infection

**DOI:** 10.1101/110304

**Authors:** Valérie Poirier, Gal Av-Gay, Yossef Av-Gay

**Author notes:** Corresponding author (YAG).

## Abstract

Alveolar macrophages serve as the first line of defence against microbial infection, yet provide a unique niche for the growth of *Mycobacterium tuberculosis*. To better understand the evasive nature of the tubercle bacilli and its molecular manifest on the macrophage response to infection, we conducted a global quantitative proteomic profile of infected macrophages. By examining four independent controlled infection experiments, we detected 42,007 peptides resulting in the characterization of 4,868 distinct proteins. Of these, we identified 845 macrophage proteins whose expression is modulated upon infection in all replicates. We showed that the macrophage’s response to *M. tuberculosis* infection includes simultaneous and concerted upregulation of selected proteins. Using a number of statistical methods, we identified 27 proteins whose expression levels are significantly regulated outside of a 90% confidence interval about the mean. These host proteins represent the macrophage transcriptional, translational, and innate immune response to infection as well as its signaling capacity. The contribution of PtpA, an *M. tuberculosis* secreted virulence factor, modulated the expression levels of 11 host macrophage proteins, as categorized by RNA metabolism, translation, and cellular respiration.

## Introduction

*Mycobacterium tuberculosis,* the etiological agent of Tuberculosis (TB), is one of the most prevalent infectious agents worldwide. An estimated 8.6 million new cases and approximately 1.3 million deaths occur annually (WHO Report, 2013) giving TB the highest mortality rate of any infectious bacteria [1]. Co-infection with HIV and the emergence of multidrug-resistant strains poses a significant challenge to treating the disease and increasing the survival rate. Thorough characterization of key physiological events of both the bacterium and the human host are needed for better understanding of the disease in order to establish new and innovative drug development programs.

Alveolar macrophages, paradoxically, serve both as first line of defence against *M. tuberculosis* and the bacterium’s natural habitat [2]. Normally, macrophages phagocytose and digest invading microorganisms leading to the initiation of an immune response [3]. This multistep process is termed phagosome maturation whereby phagosomes first acidify and then fuse with proteolytic lysosomes. However, *M. tuberculosis* circumvents the macrophage killing machinery and establishes long-term infection, enabling it to replicate within the infected macrophage. To achieve “successful” infection, *M. tuberculosis* actively interferes with host trafficking events by secreting virulence factors [3]. Among these, *M. tuberculosis* secretes the protein phosphatase Protein tyrosine phosphatase A (PtpA), which prevents macrophage acidification, the phagosome-lysosome fusion processes [4, 5] and innate immune response [6]. This allows *M. tuberculosis* to replicate and persist inside the macrophage while avoiding proteolytic degradation and antigen presentation required to initiate an adaptive immune response [2].

The intimate interaction between *M. tuberculosis* and the macrophage prompted a wide variety of genomic, transcriptomic and proteomic studies which has increased our understanding of the interface between the host macrophage and *M. tuberculosis*. In recent years, several proteomic analyses of mycobacteria and host immune cells have been conducted. The majority of these studies have investigated the mycobacterial proteome after engulfment by the macrophage [7-13], and, surprisingly, only a handful has examined the macrophage’s protein expression patterns during infection [14-18].

In this study, we analyzed the global protein expression profiles of differentiated monocytes; human macrophage-like cells (THP-1) infected with *M. tuberculosis* H37Rv, and compared them to the proteomic profiles of uninfected macrophages. Moreover, due to its role in phagosome maturation arrest and inhibition of apoptosis [4, 19, 20], we further extended our study to identifying the contribution of PtpA by comparing the proteome of macrophages infected with the PtpA-null mutant *M. tuberculosis* (Δ*ptpA*) to that of macrophages infected with the parental *M. tuberculosis* strain. To analyze the extent of proteomic reprogramming during *M. tuberculosis* infection, we used a multiplexed peptide quantitation methodology, termed Isobaric Tags for Related and Absolute Quantitation (iTRAQ). iTRAQ is a quantitative MS-based proteomic approach employed to perform a global proteomic analysis of cellular proteins [21]. Used in conjunction with tandem mass spectrometry (MS/MS) to identify the resultant peptides, iTRAQ precisely quantifies and identifies cellular proteins by placing isobaric mass labels on peptides in a peptide mixture [22].

In total, four independent iTRAQ analyses were performed and 42,007 peptides from cellular extracts of macrophages were detected resulting in the identification of 9,784 proteins, 4,868 of which are distinct proteins. Of these, 845 proteins were identified in all four replications of the experiment. We further refined the list and identified 27 proteins that are modulated in *M. tuberculosis* infection. We selected five proteins whose expression showed significant modulation by infection for quantitative PCR, transcriptional analysis, and Western blot for validation of protein expression. These include the upregulated ICAM1, IL1β, and SOD2, and the downregulated CTSG and MIF. We also analyzed consistently up- or downregulated proteins across the two independent iTRAQ analyses involving Δ*ptpA* and identified 1,281 distinct proteins, 11 of which are significantly modulated by PtpA. In brief, our analyses demonstrate that *M. tuberculosis* infection causes an upregulation of proteins involved in immunity, transcription and translation, and a downregulation of signaling proteins. Moreover, we observed a pronounced PtpA-dependent downregulation of proteins involved in RNA metabolism and cellular respiration. To the best of our knowledge this is the first study to simultaneously determine proteome-wide protein relative expression levels of human macrophage-like cells infected with *M. tuberculosis,* and outline molecular signatures of the global and PtpA-dependent proteomic pattern of macrophages during *M. tuberculosis* infection.

## Experimental Procedures

### Tissue Culture Maintenance and Differentiation

The human monocytic leukemia cell line THP-1 (TIB-202; ATCC) was cultured in RMPI 1640 medium (Sigma-Aldrich) supplemented with 10% FBS (PAA Laboratories Inc.), 1% L-glutamine, 1% penicillin and 1% streptomycin (StemCell). Cells were seeded in 10 cm (diameter) tissue culture dishes at a density of 7.0 × 10^6^ cells/dish and differentiated into a macrophage-like cell line with 20 ng/ml phorbol myristate acetate (PMA) (Sigma-Aldrich) in RPMI 1640 medium supplemented with 10% FBS and 1% L-glutamine (incomplete RPMI) at 37 °C in a humidified atmosphere of 5% CO_2_ for 18 h.

### Macrophage Infection

Bacterial cells were washed with Middlebrook 7H9 broth supplemented with 0.05% (v/v) Tween 80 (Sigma-Aldrich). Infection of THP-1 cells was performed using 10% human serum-opsonized *M. tuberculosis* at a multiplicity of infection (MOI) of 10:1 in RPMI 1640 medium. After 3 h of incubation at 37 °C and 5% CO_2_, cells were washed with RPMI 1640 medium to remove non-internalized bacteria and re-incubated at 37 °C and 5% CO_2_ in incomplete RPMI containing 100 μg/ml gentamicin (Invitrogen) for 4 h.

### Macrophage Cellular Extraction for iTRAQ Analysis

Four hours post-infection, *M. tuberculosis-ΔptpA-,* and Δ*ptpA*::ptpA-infected THP-1 cells were washed twice with RPMI 1640 medium and scraped in 3 ml of the medium. Cells were centrifuged for 10 min at 670 RPM, re-suspended in 490 μl iTRAQ lysis buffer (25 mM ammonium bicarbonate, 5 mM sodium fluoride, 5 mM sodium orthovanadate, 10 mM sodium β-glycerophosphate pentahydrate, pH 7-7.5) and transferred to screw-cap tubes containing 0.5 g of 0.5 mm glass beads. Ten μl of 10% sodium dodecyl sulphate (SDS) were added to the tubes and cellular extracts were bead-beated (BioSpec Products Inc.) for 2 min (30 sec on/30 sec off) at a speed of 5.0. The lysates were then centrifuged for 20 min at 13,000 RPM and the supernatants passed through a 0.22 μm filter column (Millipore Corporation).

### iTRAQ and LC-MS/MS Analyses

Protein concentrations of cell extracts of infected macrophages were determined using a bicinchoninic acid assay (Sigma-Aldrich). Protein samples (200 μg from each test) were sent to the University of Victoria Genome British Columbia Proteomics Centre for iTRAQ analyses. In brief, 85 μg of each sample were precipitated overnight in acetone at 4 °C and then resolubilized in 0.5 M TEAB, 0.2% SDS, Proteins were then reduced with TCEP and alkylated with MMTS. Proteins were digested in solution with trypsin (Promega) at 37 °C overnight and labeled with the appropriate iTRAQ label (113-119, 121) at room temperature for 1 h. iTRAQ labeled peptides were then combined and separated by cation exchange HPLC. HPLC fractions containing peptides were concentrated by speed-vac, then analyzed and quantified by LC-MS/MS. The length of the reverse gradient used was 2 h per HPLC fraction. Samples were analyzed by reversed phase nanoflow (300 nL/min) HPLC with nano-electrospray ionization using a LTQ-Orbitrap mass spectrometer (LTQ-Orbitrap Velos, Thermo-Fisher) operated in positive ion mode.

### Data Analysis and Interpretation

All data was analyzed using Proteome Discoverer 1.4 (Thermo Fisher Scientific) and the search engine MASCOT v2.4 (Matrix Science). Raw data files were searched against the Uniprot-SwissProt database with allspecies filter and mammalian species only.

### Statistical Analysis

Raw data files from each of the four experimental trials were collected and analyzed using R (gnu). All four trials included *M. tuberculosis* /uninfected ratios for detected proteins. Additionally, the first two trials included Δ*ptpA*/uninfected and Δp*tpA*::ptpA/uninfected ratios. Approximately two thousand proteins were detected in each trial, of which only 845 were detected in all trials. All data was log-transformed and each trial was normalized individually. Trials were normalized before elimination of contaminants and missing data.

For detection of anomaly, protein ratios of uninfected and *M. tuberculosis-infected* macrophages were compared using two methods. The first involved isolating all proteins for which the expression ratio in at least three of four assays was one standard deviation above/below the mean expression ratio. For normalized data, the mean for each assay is zero and the standard deviation is one. For instance, to isolate upregulated proteins, all proteins with a normalized log-ratio above one in at least three of the four assays were kept. The second method involved taking the mean ratio for each protein over all four assays and isolating those proteins for which this mean was outside of the ±1 standard deviation interval about the mean.

A method was implemented for the initial two iTRAQ assays which included all three treatment conditions, i.e. infection with *M. tuberculosis,* Δ*ptpA,* and Δ*ptpA*::ptpA. This was inspired by the apparent co-linearity between each of these covariates. Proteins for which the infected/uninfected ratio was 1.8 standard deviations above the mean for all three possible infected/uninfected ratios were considered as consistently upregulated proteins for that sample, and therefore not PtpA-specific. The same was done to isolate proteins consistently downregulated for each sample. These generally up- or downregulated samples had proteins ranked based on their mahalanobis distances [23] from the mean of the given sample where the three variables used in calculation of the correlation matrix were the three infected treatment conditions. Furthermore, the remaining proteins were analyzed in accordance to their *M. tuberculosis/*Δ*ptpA* ratios, organized into up- and downregulated lists based on the 1.8 standard deviation criterion, and ranked based on their distance from the mean.

A third method was used to compare protein ratios of uninfected and *M. tuberculosis-infected* macrophages as well as protein ratios of *M. tuberculosis-* and Δ*ptpA*-infected macrophages. Using the null hypothesis that the mean normalized ratio is equal to one, a series of *t* tests was implemented to isolate all proteins that rejected this null hypothesis at a *p*-value of less than 0.05. The proteins identified were shown statistically to be significantly upregulated or downregulated given the data. Some of these proteins, however, have mean ratios only marginally above or below the mean and therefore this statistical significance may not correspond to significance in a biological context.

### Macrophage Cellular Extraction for Western Blot Analysis

Four hours post-infection, infected THP-1 cells were washed twice with cold PBS and cellular extracts were harvested in lysis buffer (20 mM Tris, 5 mM EDTA, 1% Triton X-100, 1 mM DTT, 1 mM phenylmethylsulfonyl fluoride (PMSF), 1 mM phosphatase inhibitor cocktail (Sigma-Aldrich), pH 7.2) by drawing the solution in and out of a blunt syringe 15-20 times. The cellular extracts were centrifuged for 10 min at 13,000 RPM and passed through a 0.22 μm filter column.

### Western Blot Analysis

*In vivo* Western blot analyses of cell extracts harvested 4 h post-infection from infected THP-1 cells were performed as follow: 100 μg of cell extracts were resolved by SDS-PAGE and transferred onto a nitrocellulose membrane (Bio-Rad Laboratories). The blots were probed with affinity purified mouse monoclonal anti-ICAM1 (eBioscience) (final IgG dilution, 1:500), affinity purified rabbit polyclonal anti-CTSG (Antibodies-online,com) (final IgG dilution, 1:500) or affinity purified rabbit polyclonal anti-IL1β (Antibodies-online,com) (final IgG dilution, 1:1,000) and incubated overnight at 4 °C. For detection of ICAM1, Alexa Fluor 780 goat anti-mouse (Mandel Scientific) antibody was used as the secondary detection reagent (final IgG dilution, 1:5,000). For CTSG and IL1β, Alexa Fluor 680 goat anti-rabbit (Invitrogen) antibody was used (final IgG dilution, 1:10,000). An Odyssey Infrared CLx Imager (LI-COR Biosciences) was employed for chemiluminescence detection. To ensure identical protein loading of the different samples, Ponçeau staining of the blots was performed.

### Macrophage RNA Extraction and cDNA Synthesis

Total RNA was extracted from *M. tuberculosis-,* Δ*ptpA-* and Δ*ptpA::.ptpA*-infected THP-1-derived macrophages (7.0 × 10^6^ cells) at defined time points (2, 4 and 18 h) using the RNAspin Mini Kit according to the manufacturer’s instructions (GE Healthcare). RNA was reversed transcribed to cDNA using the EasyScript cDNA Synthesis Kit following the manufacturer’s protocol (ABM). For each cDNA synthesis, 1 μg of total RNA, measured by an Epoch Microplate Spectrophotometer (BioTek), and 0.5 μΜ oligo(dT) oligonucleotide primers were used.

### Quantitative Polymerase Chain Reaction (qPCR)

Primers specific for *ICAM1, IL1β, SOD2, CTSG* and *MIF* mRNA were designed using Primer-BLAST (National Center for Biotechnology Information) (S1 Table). Control PCR amplifications for the expression of the gene-specific mRNAs were performed on cDNA templates from uninfected PMA-differentiated THP-1 cells to confirm the specificity of the designed primers. Each qPCR reaction contained 2 X EvaGreen qPCR Mastermix (ABM), 15 ng cDNA and 1 μM of each primer and was analyzed in quantification mode on a DNA Engine Opticon instrument (Bio-Rad Laboratories). The following cycling conditions were used. 95 °C for 10 min, 40 cycles of 95 °C for 15 sec, 52 °C for 15 sec, and 60 °C for 30 sec with data collection during each cycle. Mock reactions (no reverse transcriptase) were also included with each experiment to confirm the absence of genomic DNA contamination. Ct values were converted to copy numbers using standard curves. Results were analyzed using GraphPad Prism 5.0 software, All values of gene-specific mRNA were internally normalized to cDNA expression levels of housekeeping gene *GAPDH.*

## Results

### Modulation of Host Protein Expression in Response to *M. tuberculosis*

To study the effect of *M. tuberculosis* on the macrophage proteome, three different strains were used for the infection experiments; the wild-type H37Rv strain (WT), the gene knockout strain Δ*ptpA* (KO), and the corresponding complemented strain (CO) where the original *ptpA* gene was reintroduced via a replicative plasmid into the mutant strain. Four independent sets of THP-1 cells infected with *M. tuberculosis* and the corresponding controls were analyzed resulting in the identification of 4,868 unique proteins and their corresponding levels in each one of the experiments. Disparity in values obtained for the expression ratios was observed between experiments (Fig. 1A); in three of the four infection experiments, infected versus uninfected log-ratios displayed a mean and a 95% confidence interval below zero. Log-ratios in the fourth experiment displayed a mean and 95% confidence interval above zero. Thus, we normalized the data so that each sample has a standard deviation of approximately one and a mean of zero (Fig. 1B). In this manner, the relative modulation of protein expression could be measured in relation to all other proteins in the same experiment, across samples and between independent experiments.

**Fig. 1.**
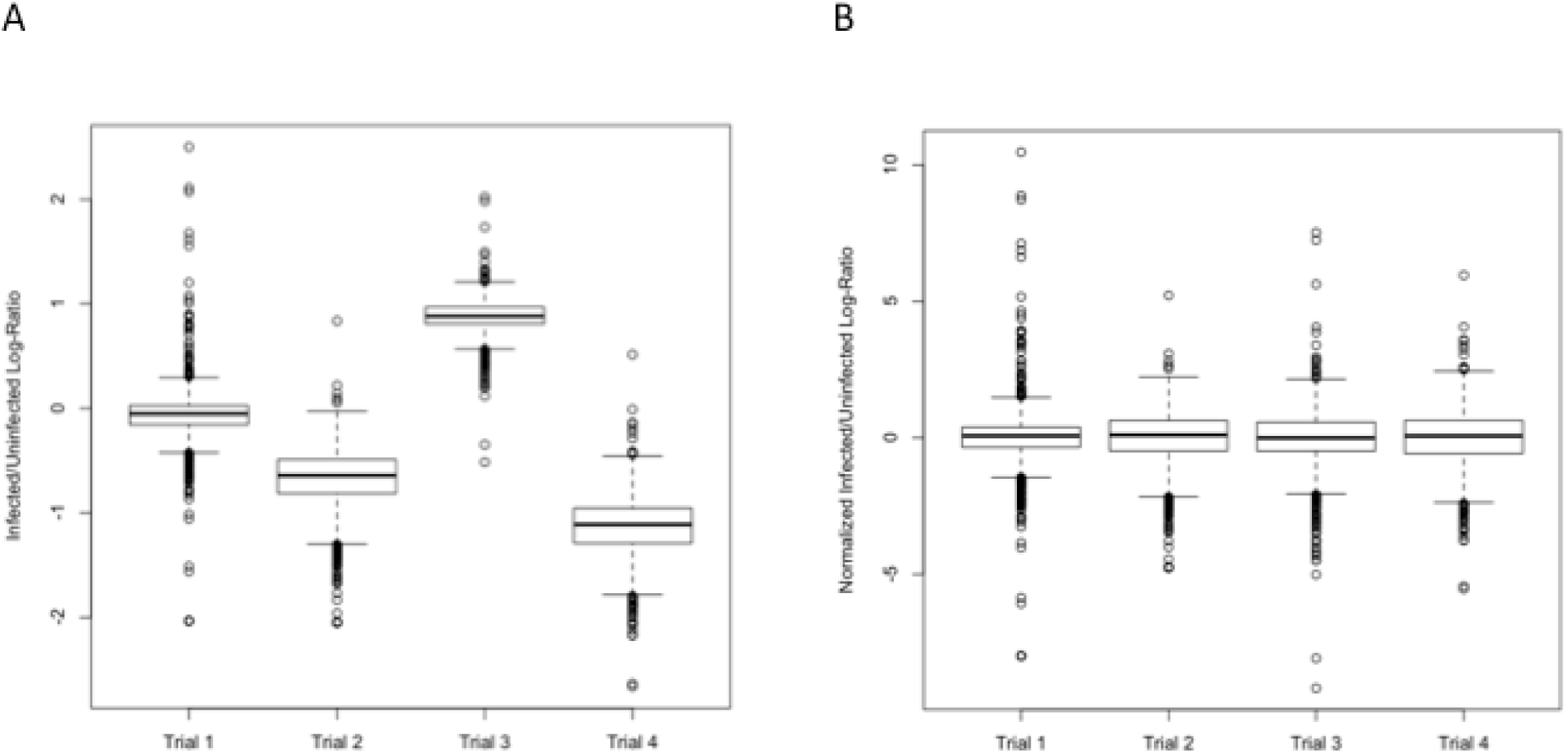
Boxplots of Host Proteins Obtained from Infected Versus Uninfected Macrophages. Log-ratios measured 4 h post-infection. (A) Log-ratios for four trials of experiments where ratios are between *M. tuberculosis-infected* and uninfected macrophages. (B) Normalized log-ratios for four trials of experiments where ratios are between *M. tuberculosis-infected* and uninfected macrophages. Each sample is normalized to have a mean and a standard deviation of approximately zero and one, respectively.

To determine the general effects of *M. tuberculosis* on host protein expression levels, we analyzed the data four ways. First, we compared log-ratios of protein expression from uninfected cells to infected ones and selected the 845 proteins present in all four independent experiments. We selected proteins for which this ratio fell outside of a one standard deviation interval about the mean for at least three of the four experiments (Table 1). The second and third approaches considered samples taken 4 and 18 h after infection using the same four independent sets of infected THP-1 cells. Proteins that exhibited a relative change in their corresponding ratios outside an interval two standard deviations above and below the mean for all three strains tested were identified and considered as general responses to *M. tuberculosis* infection independent of PtpA (S2 and S3 Tables). Lastly, the fourth approach considered all four experiments taken 4 h post-infection and used the null hypothesis that the mean normalized ratio is equal to one. A series of *t* tests was then implemented to identify all proteins that rejected this null hypothesis. We isolated all proteins for which the *p*-value was less than 0.05 and identified 52 proteins showing 95% confidence intervals above or below the mean (S1 Fig., S4 Table).

**Table 1.**
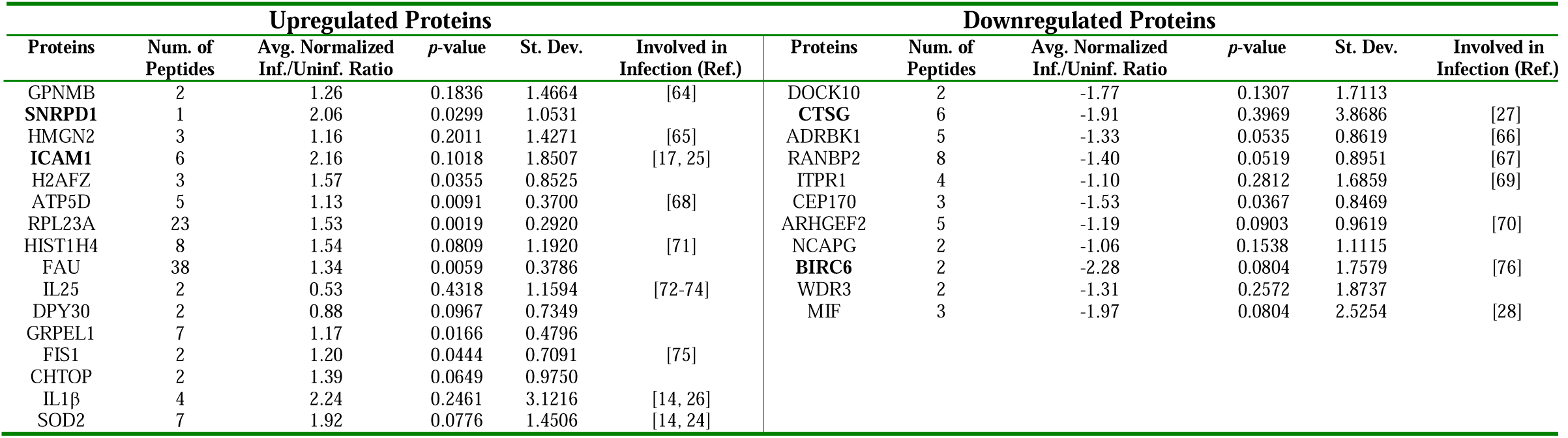
*M. tuberculosis-Infected* Macrophage Proteins Modulated One Standard Deviation Above or Below the Mean. This list was obtained by eliminating all proteins for which less than three sample ratios were within one standard deviation about the mean. Proteins in bold are proteins for which the average standardized *M. tuberculosis/uninfected* ratio (over all samples) was above or below 1.8 standard deviations about the mean. Proteins in this list are not ranked. For each protein, two-tailed *p*-values were calculated using a *t* test over the four trials. *p*-value < 0.05 indicates significance at the 95% level. *p*-value < 0.1 indicates significance at the 90% level.

In defining a robust notion of the macrophage response to infection, some data analysis difficulties were encountered. For example, one analysis approach involves classifying proteins, from infected versus uninfected macrophages, with a log-ratio above one as upregulated, and below one as downregulated. Hypothesis testing using this definition identifies a number of significant proteins, as in the fourth approach; however, exploratory analysis of our raw data suggests that many of these protein ratios were only very slightly above or below the mean. Thus, a second, more robust hypothesis testing approach was formulated where upregulated proteins are defined as those with log-ratios more than one standard deviation above the mean, and downregulated proteins are defined as those with log-ratios more than one standard deviation below the mean. Unfortunately, hypothesis testing using these definitions resulted in no tests with a significant *p*-value. As an alternative, we chose to identify all proteins for which at least three of four log ratios were more than one standard deviation above the mean, or one standard deviation below the mean, but not both. Using this approach, we can get as close as possible to isolating proteins that can be classified as significantly up- or downregulated for this specific data set.

Furthermore, to identify the expression trends of host proteins modulated by *M. tuberculosis* infection, we selected proteins that are either consistently up- or downregulated for each of the strains and determined their expression levels by the ratio between infected over uninfected macrophages. The ratios were plotted against one another for each individual sample. We observed that upregulated proteins show a strong linear relationship between each ratio (Figs. 2A-B), indicating that upon independent *M. tuberculosis* infection, similar proteins become upregulated at similar rates. Alternatively, there appears to be a lack of linearity of the macrophage response in terms of downregulating protein levels (Figs. 2A-B, inserts).

**Fig. 2.**
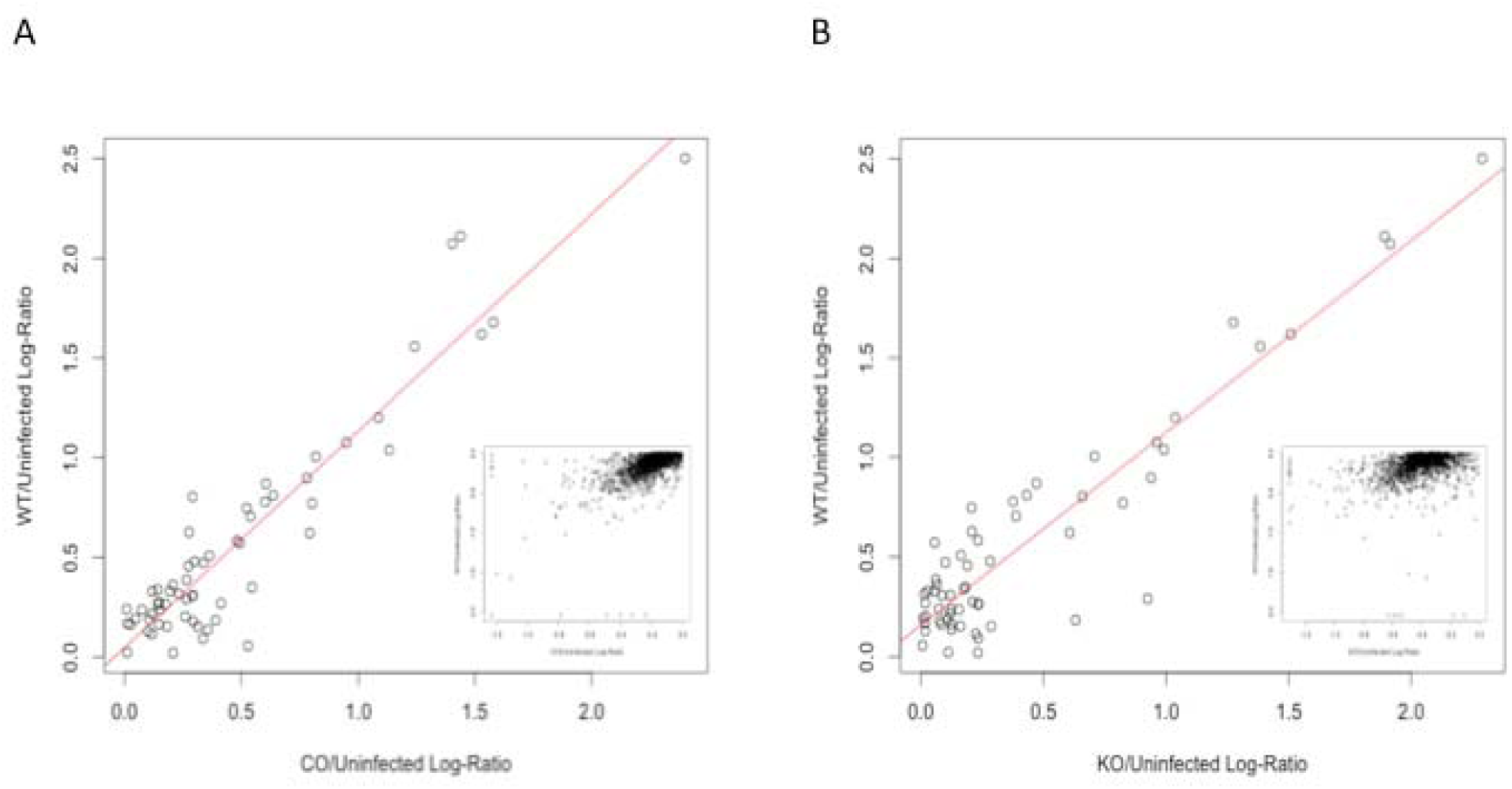
*M. tuberculosis-Modulation* of Host Protein Expression. Proteins for which the infected to uninfected expression ratios are above and below the mean 4 h post-infection. (A) The *M. tuberculosis/*Δ*ptpA: ptpA* log-ratio is plotted and shows a strong linear relationship between these expression ratios for upregulated proteins. The insert represents the plotted *M. tuberculosis/*Δ*ptpA::ptpA* log-ratio and shows no linear relationship between these expression ratios for downregulated proteins. (B) The *M. tuberculosis/*Δ*sptpA* log-ratio is plotted and shows a strong linear relationship between these expression ratios for upregulated proteins. The insert represents the plotted *M. tuberculosis/*Δ*ptpA* log-ratio and shows no linear relationship between these expression ratios for downregulated proteins.

### Identification of Macrophage Proteins Modulated by *M. tuberculosis* Infection

To identify the statistically relevant host proteins modulated by *M. tuberculosis* infection, we selected from the 845 distinct proteins those whose average expression level in infected over uninfected ratios fell outside of ±1 standard deviation from the global mean for at least three of the four experiments (Table 1). We further shortlisted proteins by taking the mean ratio for each protein over all four trials and excluded those whose ratios were within 1.8 standard deviations of the mean (Table 1). Using these two methods, we identified 27 unique host proteins to be modulated in *M. tuberculosis* infection (Fig. 3, Table 1).

**Fig. 3.**
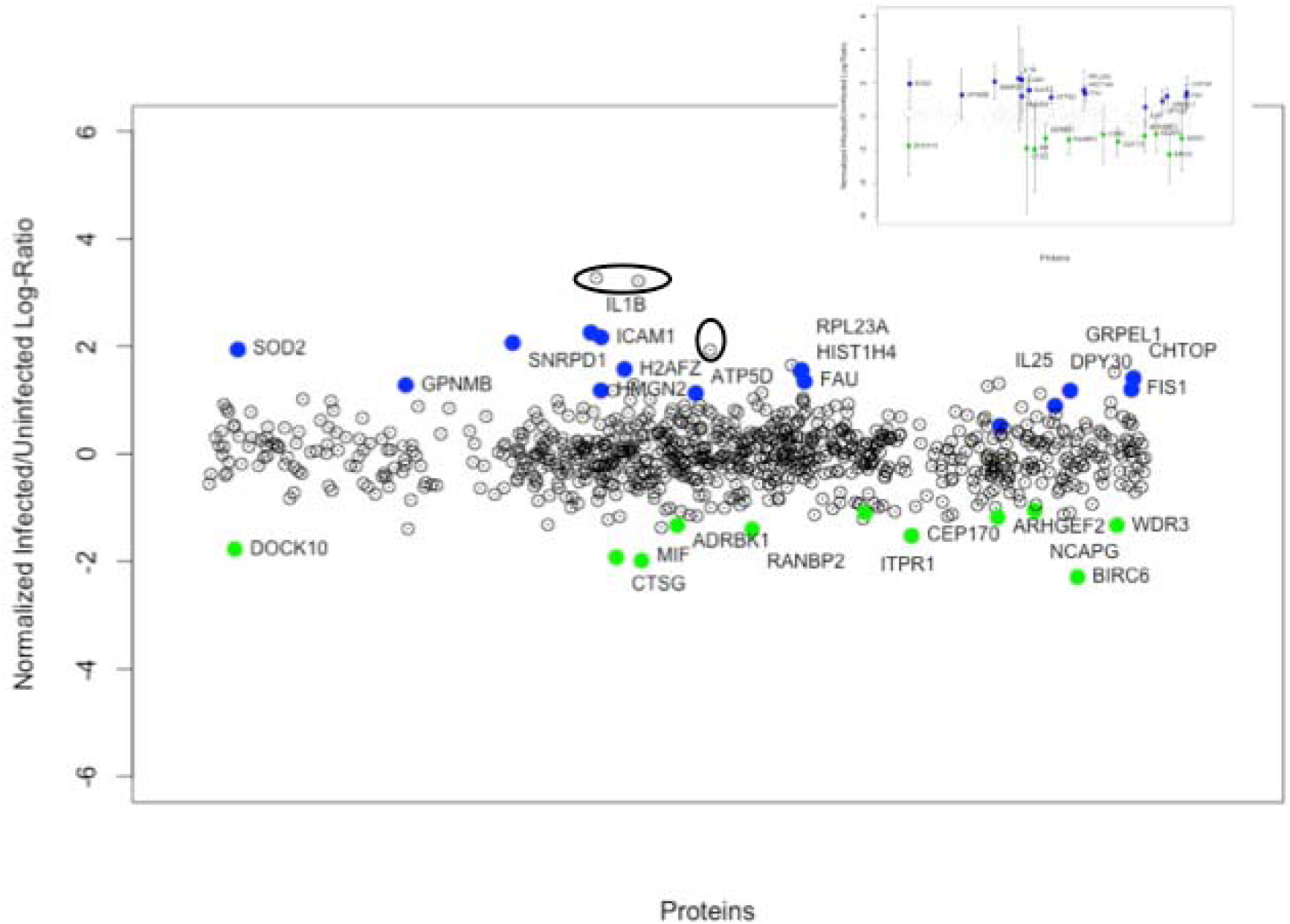
Global Expression Level Scatterplot of Normalized Infected Versus Uninfected Log-Ratios. Includes 845 proteins detected in all four trials where highlighted proteins are those with mean expression ratios above or below a one standard deviation confidence interval about the mean. Blue dots correspond to upregulated proteins, and green dots correspond to downregulated protein, as per Table 1. The extreme values marked by a circle were determined to be contaminants (P35527, P04264, P13645).

Sixteen of the identified proteins were previously reported to be involved in the macrophage response to microbial infections. For instance, upregulated SOD2 [14, 24], ICAM1 [17, 25], and IL1β [14, 26], and downregulated CTSG [27] and MIF [28] have all been involved in the innate immune response to pathogens. Conversely, among the 11 proteins that have previously never shown to be associated with or affected by infection, we identified the upregulated RNA metabolism protein SNRPD1, the chromatin remodeling and transcription protein H2AFZ, the ribosomal protein RPL23A, and the downregulated signaling protein DOCK10.

### Validation of Proteomics Expression Levels

To experimentally examine the validity of our findings, we selected five representative proteins from those listed in Table 1 and compared our iTRAQ findings to those obtained from either Western blot analysis (for protein levels) or quantitative PCR (qPCR for transcription levels). We selected five proteins previously reported to be modulated in microbial infections.

Western analysis was conducted for ICAM1, IL1β and CTSG. Based on the iTRAQ results (Table 1), protein levels of ICAM1 and IL1β were expected to be upregulated in infected cells and those of CTSG, to be downregulated. As anticipated, levels of ICAM1 and IL1β in *M. tuberculosis*-infected cells were upregulated (Figs. 4A-B) correlating with and confirming the modulations observed in our proteomic analyses. However, as seen in Fig. 4C, in contrast to our iTRAQ findings, CTSG levels remained unchanged when comparing infected to uninfected macrophages.

**Fig. 4.**
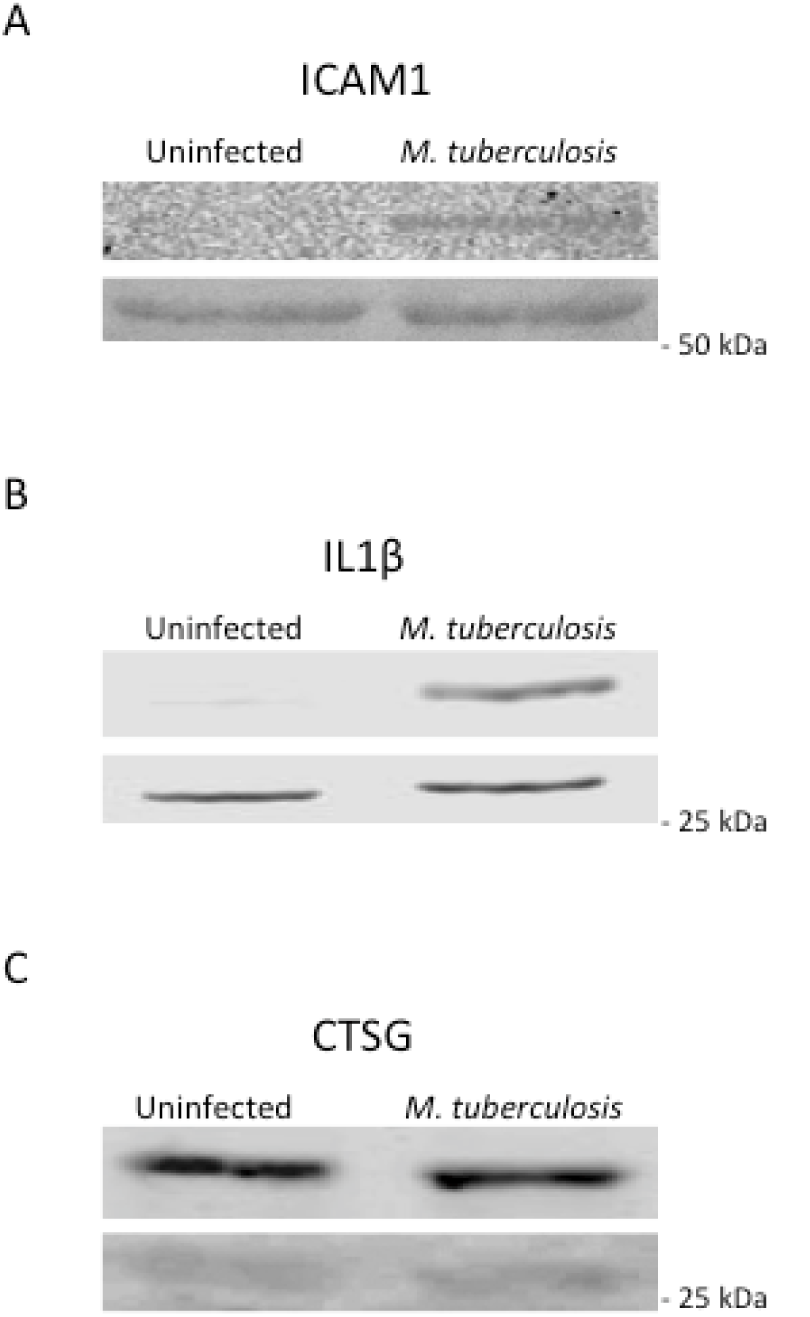
Western Analyses of *M. tuberculosis-Dependent* Host Protein Expression Levels. THP-1 cells were infected with *M. tuberculosis* and cellular extracts were harvested 4 h post-infection. A total of 100 μg of cellular extract were employed for Western blotting in which (A) anti-ICAM1, (B) anti-IL1β, and (C) anti-CTSG antibodies were utilized. The molecular mass of ICAM1 is 57.825 kDa, of IL1β is 30.748 kDa, and of CTSG is 28.837 kDa. The bottom panels represent the Ponçeau stained membrane showing equal loading of samples.

The mRNA levels of *ICAM1, IL1β SOD2, CTSG,* and *MIF* were quantified by qPCR. The average mRNA fold change was assessed 2, 4 and 18 h post-infection and was compared to the relative change in protein levels observed in the corresponding iTRAQ experiments. As seen in Fig. 5, qPCR analysis revealed upregulation in mRNA levels of *ICAM1, IL1β* and *SOD2* in cells infected with *M. tuberculosis* 4 h post-infection (Figs. 5A-C). The increase in mRNA abundance correlates with our proteomic data (Table 1). Transcription levels of *CTSG* and *MIF* in infected cells were unchanged, contradicting the proteomic results that detected downregulation 4 h post-infection (Figs. 5D-E, Table 1). Levels of *CTSG* mRNA remained unchanged in infected macrophages and increased 18 h post-infection (Fig. 5D), while mRNA levels of *MIF* showed no significant change in any of the time points monitored (Fig. 5E).

**Fig. 5.**
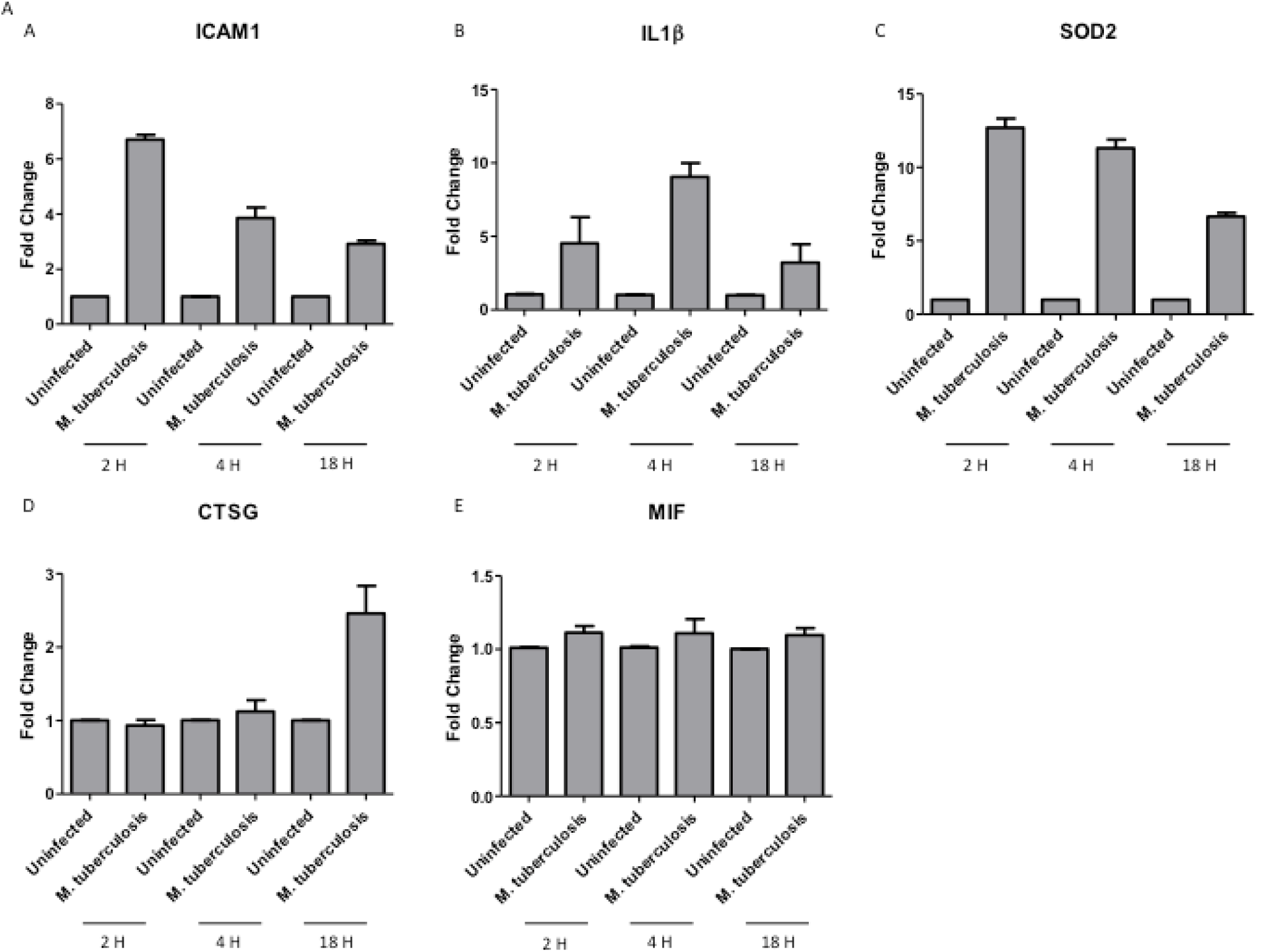
Transcriptional Levels of Five *M. tuberculosis-Modulated* Proteins. Quantitative PCR analysis comparing mRNA levels of (A) *ICAM1,* (B) *IL1β*, (C) *SOD2,* (D) *CTSG* and (E) *MIF* from uninfected and infected cells. RNA from uninfected and *M. tuberculosis-infected* THP-1 cells was extracted 2, 4 and 18 h after infection and reverse transcribed. Data shows the expression levels of all genes. Transcript abundance was determined relative to the housekeeping gene *GAPDH.* Data shown is the means ± standard deviation of three independent experiments.

### PtpA Effect on Macrophage Protein Expression Levels

To assess the impact of the virulence factor PtpA on the macrophage’s response to infection, we compared the macrophage proteome in response to infection with WT or KO strains 4 h post-infection and normalized the data as described above to give each sample a mean of zero and a standard deviation of approximately one (Figs. 6A-B). In total, we identified 1,281 proteins, including 11 unique macrophage proteins whose expression ratios fell outside of a ±1.8 standard deviation interval for macrophages infected with the KO strain (Fig. 7, Table 2). We chose 1.8 as a compromise because one standard deviation isn’t enough of a threshold when there are only two samples, and two standard deviations resulted in no proteins being defined as significant. Among these PtpA-dependent proteins, three have previously been reported to be involved in the infection process; the upregulated stress response protein CIRBP [29], and the downregulated signaling protein GSK3α [30], and the mitochondrial peptide transporter ABCB10 [31]. GSK3α is the only PtpA-dependent protein that has been identified before in mycobacterial infections [20, 32-34]. Among the eight other proteins, the most modulated ones include the downregulated RNA metabolism protein PRPF31, cellular respiration proteins COX7B and ATP5I, and the protein synthesis protein SARS2.

**Fig. 6.**
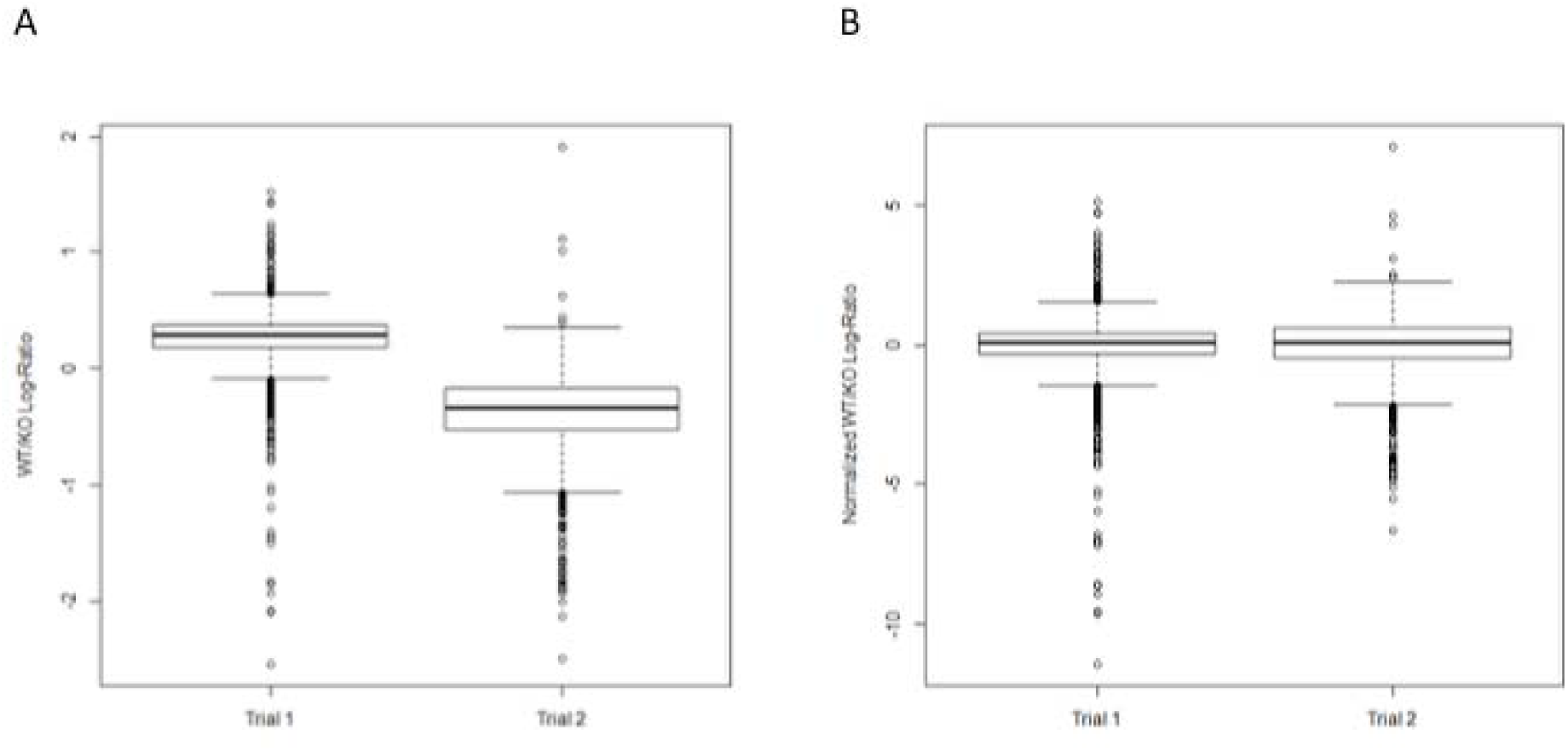
Boxplots of Host Proteins Obtained from Infected Versus Uninfected Macrophages. Log-ratios measured 4 h post-infection. (A) Log-ratios for two trials of experiment where ratios are between *M. tuberculosis-* and Δ*ptpA*-infected macrophages. (B) Normalized log-ratios for two trials of experiment where ratios are between *M. tuberculosis-* and Δ*ptpA*-infected macrophages. Each sample is normalized to have a mean and a standard deviation of approximately zero and one respectively.

**Fig. 7.**
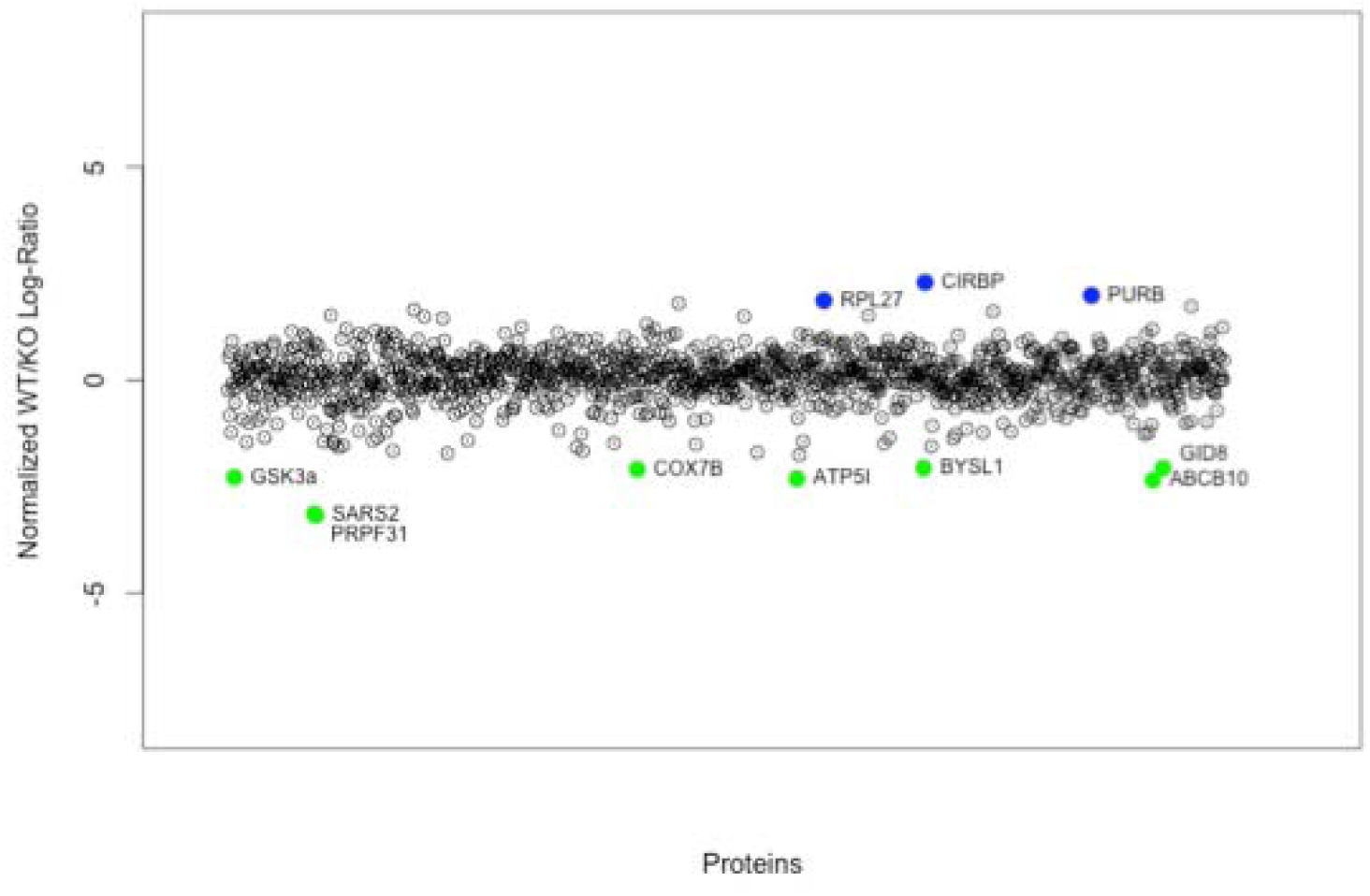
Global Expression Level Scatterplot of Normalized *M. tuberculosis* Versus Δ*ptpA* Log-Ratios. Includes 1281 proteins detected in two trials of the experiment. Highlighted proteins are those with mean *M. tuberculosis/ΔptpA* ratios above or below a 1.8 standard deviation confidence interval about the mean. Blue dots correspond to upregulated proteins, and green dots correspond to downregulated proteins, as per Table 2.

**Table 2.**
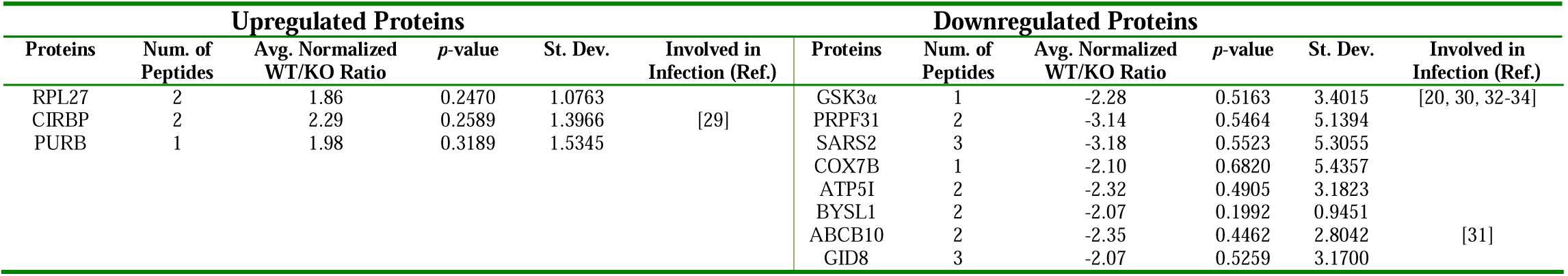
Δ*ptpA-Infected* Macrophage Proteins Modulated 1.8 Standard Deviations Above or Below the Mean. The list was obtained using two methods: (i) eliminating all proteins for which the *M. tuberculosis/*Δ*ptpA* sample ratios were within 1.8 standard deviations about the mean for both samples, and (ii) eliminating all proteins for which the mean *M. tuberculosis/*Δ*ptpA* ratio was within the ±1.8 standard deviation interval about the mean. The proteins in this list are not ranked. For each protein, two-tailed *p*-values were calculated using a *t* test over the two trials. *p*-value < 0.05 indicates significance at the 95% level. *p*-value < 0.1 indicates

Three additional lists of PtpA-dependent induced host proteins were generated using four iTRAQ trials that included all three strains of *M. tuberculosis*. This method considered two samples taken 4 and 18 h after infection. All proteins that displayed consistent behaviour (up- or downregulation) across all treatments (WT/uninfected, KO/uninfected, and CO/uninfected) were eliminated. The remaining proteins were ranked based on their WT/KO ratios. We obtained two lists of proteins that significantly exhibit PtpA-dependent behaviour. These include proteins that display higher rates (S5 Table) or lower rates (S6 Table) of expression in *M. tuberculosis-infected* macrophages compared to Δ*ptpA*-infected macrophages. The third list of PtpA-dependent host proteins was generated using the two iTRAQ experiments taken 4 h post-infection that included the WT/uninfected and the KO/uninfected ratios. With our null hypothesis being that the mean normalized expression ratio was equal to one, we identified proteins for which the *p*-value of a *t* test using the WT/KO expression ratios is less than 0.05. These proteins were selected and considered as PtpA-dependent. We isolated 58 proteins showing 95% confidence intervals above or below the mean using this method (S7 Table). However, as previously explained, analysis of significant proteins identified using this hypothesis suggests that many of these protein ratios were only very slightly above or below the mean.

## Discussion

*M. tuberculosis* has adapted well to the human host, using multiple strategies to subvert recognition and immune responses [6]. In recent years, several large-scale expression studies investigating global transcriptional responses of macrophages exposed to *M. tuberculosis* have been performed [14, 35-48]. These studies have undeniably contributed to our understanding of the extent of transcriptomic reprogramming the macrophage undergoes during mycobacterial infection. However, a broader systems-biology approach would provide a better understanding of TB specific host-pathogen interactions and macrophage biology. Furthermore, understanding the mechanisms by which *M. tuberculosis* manipulates the host response will provide fertile ground for novel interventions by means of chemotherapy or vaccine development, especially by targeting the most consistent protein changes in infected macrophages. Proteomics provides us with a higher level of understanding of biological functions than transcriptomic analysis, as it takes into account protein levels, post-translational modifications and stability, as well as protein subcellular localization.

In this study, we provide insight into the macrophage’s global response to *M. tuberculosis* infection by using a quantitative proteomic approach, and show, for the first time, how the specific virulence factor, PtpA, affects the macrophage proteome. Our first finding indicates that the macrophage acts by modulating the protein expression of a variety of specific “responder proteins” that are activated in a concerted upregulating manner by all strains examined in the study. Contrary to this, downregulation of proteins seems to be random and varies from strain to strain and between experiments. Retrospectively, it seems more logical, from an energetic point of view, to induce a well defined set of proteins required to cope with infection rather than shutting down specific sets of housekeeping or resting state homeostasis proteins which will in turn affect the macrophage’s ability to sustain infection.

### Host Proteins Targeted by *M. tuberculosis*

Using the iTRAQ methodology, we identified 845 distinct host proteins whose expression level was modified by *M. tuberculosis* infection. Among these, 27 statistically validated modulated proteins seem to form the “responder” family of proteins. These proteins were classified into functional classes (Fig. 8). Among the 16 upregulated proteins, the chromatin remodeling and transcription regulation (HMGN2, HIST1H4, H2AFZ, DPY30), transport (GPNMB, GRPEL1, FIS1), RNA metabolism (SNRPD1, CHTOP), protein synthesis and metabolism (FAU, RPL23A), and immunity and defence (IL25, IL1β) functional classes contain the most proteins. The stress response (SOD2), cellular respiration (ATP5D), and cell adhesion (ICAM1) classes are only represented by one protein (Fig. 8A, Table 1).

**Fig. 8.**
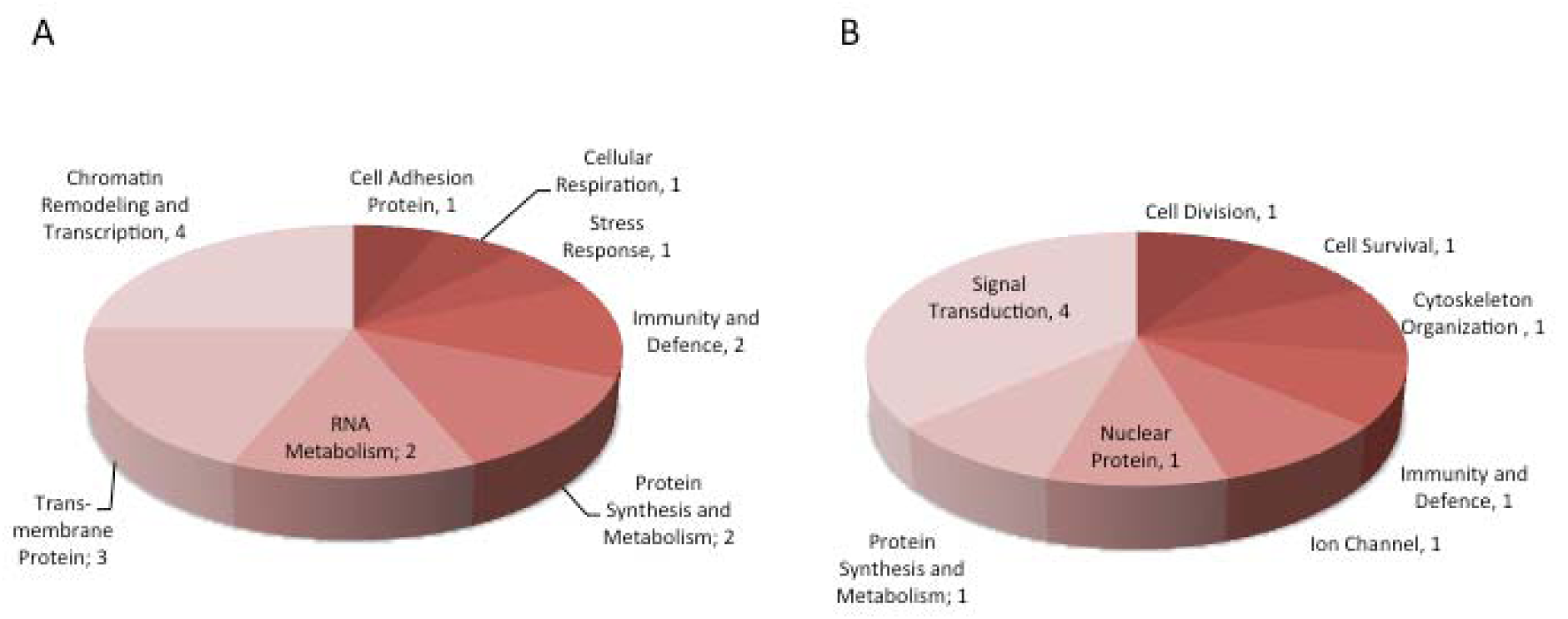
Host Functional Classes Impacted by *M. tuberculosis* Infection. The 27 proteins modulated by *M. tuberculosis* are represented as functional classes. (A) Includes the 16 upregulated proteins. (B) Includes the 11 downregulated proteins.

The upregulation of proteins involved in RNA metabolism, chromatin remodeling and transcription regulation, and in protein synthesis and metabolism, indicates that upon infection the host turns the transcriptional and translational machineries on to synthesize proteins to fight the pathogen. This is supported by the lack of downregulation of proteins involved in transcription and translation, as the majority of downregulated proteins belong to the signal transduction functional class (DOCK10, ADRBK1, RANBP2, ARHGEF2). The remaining proteins are from the protein synthesis and metabolism (CTSG), nuclear (WDR3), ion transport (ITPR1), immunity and defence (MIF), cytoskeleton organization (CEP170), cell survival (BIRC6), and cell division (NCAPG) functional classes (Fig. 8B, Table 1). The pronounced downregulation of proteins involved in signal transduction may be due to the action of virulence factors [6] secreted by *M. tuberculosis* that directly or indirectly target these host proteins, or due to an autocrine or paracrine effect which downregulates expression of signaling proteins.

### Validation of Modulation of Proteins Involved in the Innate Immune Response

To validate our TRAQ results, we investigated the correlation between the transcriptional and translational levels of five proteins whose modulation has previously been reported in mycobacterial infections. We selected SOD2 (relative fold change of 1.92) [14, 24], ICAM1 (2.16) [25], IL1β (2.24) [14, 26], CTSG (-1.91) [27], and MIF (-1.97) [28] for further analysis (Table 1).

The proteomic data of the five modulated proteins was confirmed at the transcriptional level by performing gene expression profiling using qPCR. We also verified the protein expression levels of ICAM1, IL1β and CTSG by Western blotting. As observed in the iTRAQ data (Table 1), mRNA levels of *ICAM1, IL1β* and *SOD2* were upregulated 4 h post-infection (Figs. 5A-C). The upregulated expression observed between uninfected and *M. tuberculosis*-infected cells is extremely convincing.

*Intercellular adhesion molecule-1 (ICAM1)* mRNA exhibited a four-fold increase 4 h after infection (Fig. 5A). ICAM1 is a glycoprotein expressed at the surface of immune cells that plays a role in the migration of macrophages to the site of infection and the presentation of antigens to T cells [25]. Its expression has been reported to be significantly enhanced on macrophages harbouring *M. tuberculosis* [49]. Western blot analysis of ICAM1 showed upregulated expression levels in *M. tuberculosis*-infected cells (Fig. 4A), which is consistent with our iTRAQ results (relative fold change of 2.16) (Table 1) and reports in the literature.

The cytokine *interleukin-1 beta* (*IL1β*) displayed a 10-fold increase in mRNA levels between uninfected and infected cells 4 h post-infection (Fig. 5B). IL1β is produced by activated macrophages and acts as an important mediator of the inflammatory response through its involvement in several cellular activities such as; T cell proliferation, B cell maturation and proliferation, cell differentiation, and apoptosis [26]. IL1β was previously observed in the supernatants of macrophages stimulated with *M. tuberculosis* and it is believed to play a role in the resistance of macrophages to *M. tuberculosis* infections [50]. Western blot analysis of IL1β showed a pronounced increase in expression levels in *M. tuberculosis-infected* macrophages (Fig. 4B). Protein and mRNA expression levels of IL1β were upregulated across both analytical platforms validating the iTRAQ results (relative fold change of 2.24) and reinforcing the affirmation that IL1β is an important player in the fight against *M. tuberculosis* (Table 1).

The mRNA levels of the stress response antioxidant *superoxide dismutase-2 (SOD2)* increased 12-fold 4 h post-infection (Fig. 5C) which corresponds to the upregulated protein expression observed in the iTRAQ data (relative fold change of 1.92) (Table 1). This upregulation is justified by the sudden oxidative bursts generated by the macrophage upon phagocytosis of *M. tuberculosis*. Expression of SOD2 must be increased to defend the host from its own toxic assaults from superoxide anions (O_2_^•-^) and reactive oxygen species (ROS) [51-53]. In such instances, SOD2 scavenges O_2_^•-^ offering the host cytotoxic protection. Upregulation of SOD2 expression is, therefore, a crucial defence mechanism employed by the macrophage, but also a tool used by *M. tuberculosis* to protect itself from O_2_^•-^ killing since the protection of the host from O_2_^•-^ and ROS converts the macrophage into a sanctuary for *M. tuberculosis* to grow.

The gene expression levels of *cathepsin G (CTSG)* do not correlate with its protein expression levels. As seen in Fig. 5D, mRNA levels of *CTSG* from *M. tuberculosis-infected* cells show minimal upregulation 4 h post-infection, which does not correlate with the downregulated protein levels identified by iTRAQ 4 h after infection (relative fold change of −1.91) (Table 1). Moreover, results from Western blot analysis show no change in protein expression levels of CTSG, which seemingly contradict the iTRAQ data (Fig. 4C). Transcription levels of *CTSG,* which encodes a serine protease that possesses antibacterial activities in the killing and digestion of engulfed pathogens, were shown to be downregulated in macrophages upon *M. tuberculosis* infection [27]. This is in agreement with our iTRAQ results. However, our Western blot and qPCR data does not compare with our iTRAQ results or the literature as expression levels of *CTSG* are unchanged (Fig. 4C) or slightly above basal levels (Fig. 5D). Moreover, a 2.5 fold increase in mRNA expression levels is even observed in infected cells 18 h postinfection (Fig. 5D). These findings may indicate that *M. tuberculosis* is only marginally capable of inhibiting *CTSG* expression to basal levels early during infection.

A lack of association also exists between gene and protein expression levels of migration inhibitory factor (MIF). Our qPCR data did not correlate with the proteomic results (relative fold change of −1.97) (Table 1) as a slight *MIF* upregulation is noticed in infected cells at all time points observed (Fig. 5E). MIF is a proinflammatory cytokine involved in the innate immune response to pathogens and has been shown to be secreted by and upregulated in macrophages upon exposure to *M. tuberculosis* [54]. MIF-deficient macrophages have revealed impaired mycobacterial killing [28].

The interpretation of transcriptomic data for *CTSG* and *MIF* can be variable due to the high frequency of false positives [55] and to the level of transcription of a gene not reflecting its level of expression into a protein. Indeed, mRNA transcripts may not always be translated into proteins, may be degraded [56] or, through alternative splicing, may give rise to several proteins [57]. It is also possible that, in the case of CTSG, the protease is more stable within the macrophage by means of possessing a lower degradation rate or higher half-life during infection. By contrast, *MIF* mRNA may be regulated post-translationally by events such as protein-targeted degradation. Post-translational modifications are strategies commonly employed by pathogens to modulate key host signaling pathways to their advantage. This would play a pivotal role in the pathogenesis of infection due to MIF’s important functions in immunity, and explain why downregulated protein levels were observed in the iTRAQ analyses, but not in the qPCR assay.

### Host Proteins Targeted by *M. tuberculosis* PtpA

The proteomic response of macrophages infected with the PtpA mutant strain resembles that of macrophages infected with the parental or complement strains (Fig. 2A). As observed in Fig. 2B, a strong linear relationship exists between each of the ratios indicating that upon *M. tuberculosis* and Δ*ptpA* infection, similar proteins become upregulated at comparable rates. However, little concordance is observed in the rates of protein downregulation, and, as for the other *M. tuberculosis* strains, infection with the PtpA mutant caused proteins to be downregulated at seemingly random rates (Fig. 2B, insert). By examining two independent controlled infections, we were able to identify modulation in the expression of 11 PtpA-dependent host proteins with statistical confidence (Table 2). Surprisingly, the majority of the PtpA-dependent proteins were downregulated and include members of the RNA metabolism (PRPF31, BYSL1) and cellular respiration (COX7B, ATP5I) functional classes. The other downregulated proteins belong to the transport (ABCB10), signal transduction (GSK3α), protein synthesis and metabolism (SARS2), and nuclear (TWA1) classes (Fig. 9B). The upregulated proteins are from the chromatin remodeling and transcription regulation (PURB), stress response (CIRBP), and protein synthesis and metabolism (RPL27) functional classes (Fig. 9A).

**Fig. 9.**
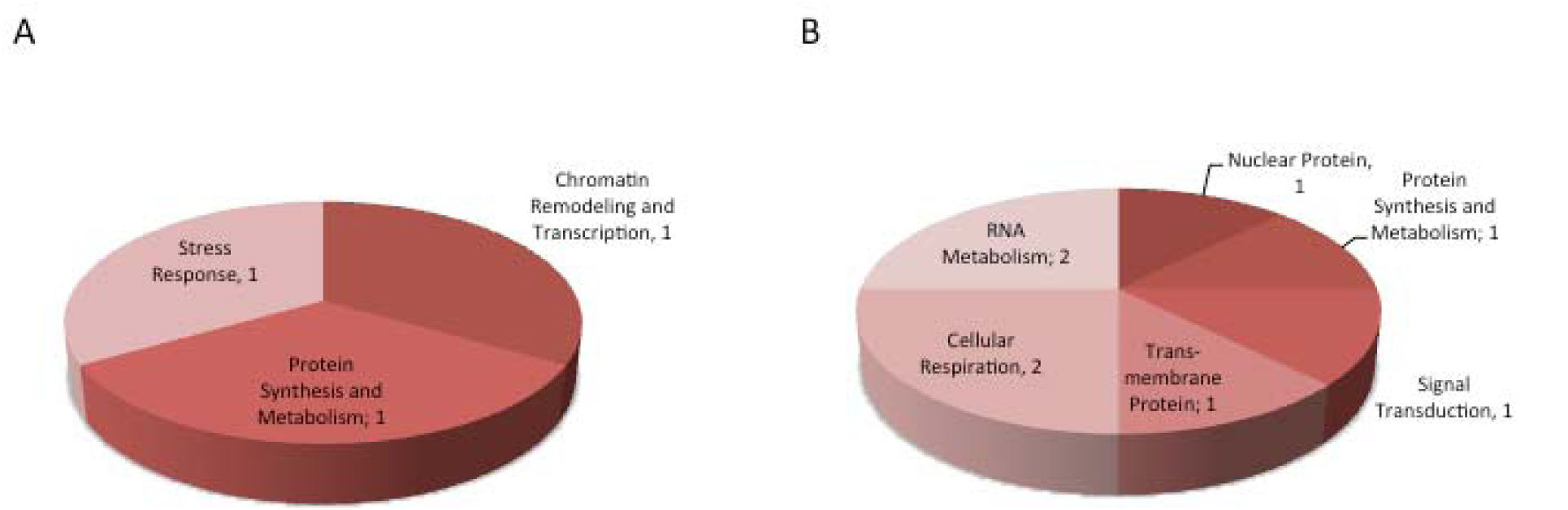
Host Functional Classes Impacted by *ΔptpA* Infection. The 11 proteins modulated by Δ*ptpA* are represented as functional classes. (A) Includes the three upregulated proteins. (B) Includes the eight downregulated proteins.

The PtpA-dependent downregulation of host proteins involved in protein synthesis and cellular respiration suggests that *M. tuberculosis* reduces the capacity of the host to produce ATP. On the one hand, this may be a survival strategy employed by *M. tuberculosis* as ATP depletion leads to necrosis of the host [58]. Indeed, certain pathogens, such as *Salmonella typhi* [59] and *Shigella flexneri* [60], prioritize host cell death by necrosis over apoptosis as apoptosis prevents pathogen propagation to neighbouring cells [61]. In addition, we have previously shown that PtpA inhibits apoptosis of the host early in infection by inhibiting activation of the apoptosis executioner, caspase-3 [20], which could indicate that *M. tuberculosis* specifically initiates necrosis early in infection for its propagation. On the other hand, previous studies have suggested that programmed necrosis of the host can serve as a backup mechanism when apoptosis is inhibited [62]. In this situation, necrosis may be the only solution considered by the host to initiate a strong inflammatory response as necrotic cells are more efficient than apoptotic cells at stimulating antigen-presenting cells and T-cell responses [63]. Thus, the impaired ATP production suggested may either be a result of *M. tuberculosis* specifically targeting energy production of the host to induce necrosis for spreading, or a voluntary action taken by the distressed host.

Previous studies from this lab analyzed the macrophage’s phosphoproteome response to infection. We investigated the phosphorylation status of key host macrophage kinases during infection with live or heat killed *M. bovis* BCG [34] or *M. tuberculosis* or Δ*ptpA* [20]. Overall, we noted changes in host signaling pathways caused by mycobacterial infection, including apoptosis (upregulation of c-JUN, SAPK and GSK3β), cytoskeletal arrangement (upregulation of α-adducin), Ca^2^+ signaling (upregulation of NR1) and macrophage activation (downregulation of PKCε) [34]. We also showed that *M. tuberculosis* PtpA dephosphorylates GSK3α as well as RAF1, PKCδ, GSK3β, PKCα/β2, and MSK1 [20]. Moreover, *in vitro* phosphorylation analysis of THP-1 proteins by recombinant PtpA resulted in the identification of multiple host signaling substrates including hVPS33B, a cognate PtpA substrate we have identified before [4, 20]. These events may contribute to downstream effects resulting in the observed protein expression we describe in this report. Future experiments will reveal the mechanisms enabling *M. tuberculosis* to invade the macrophage and the macrophage multiphasic response that enables the parasitic nature of *M. tuberculosis* infection.

## Acknowledgements

Funding for this research was provided by the Canadian Institute of Health Research (CIHR) operating grant MOP-106622 and the British Columbia Proteomics Network (BCPN). We thank Jeffrey Helm for proof reading of our manuscript, the University of Victoria Genome Proteomics Centre for their mass spectrometry analysis, and the British Columbia Centre for Disease Control for the use of the containment level 3 facility.

